# Temporal changes in surface tension guide the accurate asymmetric division of Arabidopsis zygotes

**DOI:** 10.1101/2024.08.07.605794

**Authors:** Zichen Kang, Sakumi Nakagawa, Hikari Matsumoto, Yukitaka Ishimoto, Tomonobu Nonoyama, Yuga Hanaki, Satoru Tsugawa, Minako Ueda

## Abstract

In most plants, the zygote divides asymmetrically to define the body axis. In *Arabidopsis thaliana*, the zygote undergoes polar elongation maintaining a transverse band of cortical microtubules (MTs), and divides asymmetrically forming another MT band, preprophase band (PPB). How the MT band is maintained at the actively growing cell tip and whether it contributes to PPB formation remain elusive. By combining live-cell imaging and mechanical simulation, we show that zygote elongation induces a temporal change (large material derivative) in surface tension at the growing tip to maintain the MT band, which in turn supports polar elongation. The MT band then guides PPB to determine the cell division site. Therefore, autonomous mechanical feedback between cell elongation and MT organization ensures the zygote division asymmetry.

## Introduction

In most plants, a unicellular zygote produces the multicellular plant body; this begins with the zygotic asymmetric cell division that prefigures the apical–basal axis. In *Arabidopsis thaliana* (Arabidopsis), the zygote undergoes polar elongation before dividing asymmetrically and thus the apical–basal direction is already evident during the polar elongation (*1*). Based on live-cell imaging and image quantification, we recently showed that the zygote tip, where the sperm cell fused to the egg cell, grows towards apical (*2*). In polar elongating plant cells such as root hairs and pollen tubes, actin filaments (F-actin) and microtubules (MTs) align longitudinally along the direction of cell growth (*3, 4*). Zygotes have longitudinally aligned F-actin but form a transverse band of cortical MTs at the subapical region of the growing tip (*5*). Although this subapical MT band is retained at the actively renewing zygote tip, the underlying mechanism and its role are unclear.

In addition, plants form a preprophase band (PPB) of MTs at the cortical region surrounding the nucleus to guide the formation of cell division plane (*6*). In asymmetrically dividing Arabidopsis cells, such as initial cells of stomata and lateral roots, the nucleus polarly migrates and the division plane forms at its nucleus final position (*7, 8*). In the zygote, the nucleus migrates apically along F-actin, and the PPB and cell division plane form at the apical region (*5*), but the relationship between these structures and the subapical MT band is unknown.

## Results

### The MT band is maintained at a constant distance from the apical cell pole during zygote elongation

We analyzed the dynamics of zygote growth and MT structure in Arabidopsis based on time-lapse observations using MT/nucleus markers and two-photon excitation microscopy (2PEM) (Fig. 1A and Movie S1). As shown previously, we detected the subapical MT band during zygote elongation (*5*). The MTs then form the mitotic apparatus, including the PPB, spindle, and phragmoplast, leading to asymmetric cell division (see below). We performed quantitative analysis of the images to capture the spatiotemporal dynamics of cell growth, such as cell contour, cell length (*L*), and tip radius (*r*_*e*_), as well as detecting the MT band based on truncated Gaussian fitting of MT signal distribution along the cell centerline (Fig. 1B, Fig. S1, and Movie S1, see Materials and Methods). As previously observed, the zygote entered the rapid growth stage (RGS), with its growth rate (*dL/dt*) transiently increasing after a gradual decrease in cell tip radius (Fig. 1C and D) (*2*). The subapical MT band remained at a constant position relative to the cell tip during zygote elongation prior to the formation of the mitotic apparatus (Fig. 1E). These features were observed in all samples examined, although individual zygotes differed in cell growth rate and timing of the RGS (Fig. S2A-F).

**Figure 1.**
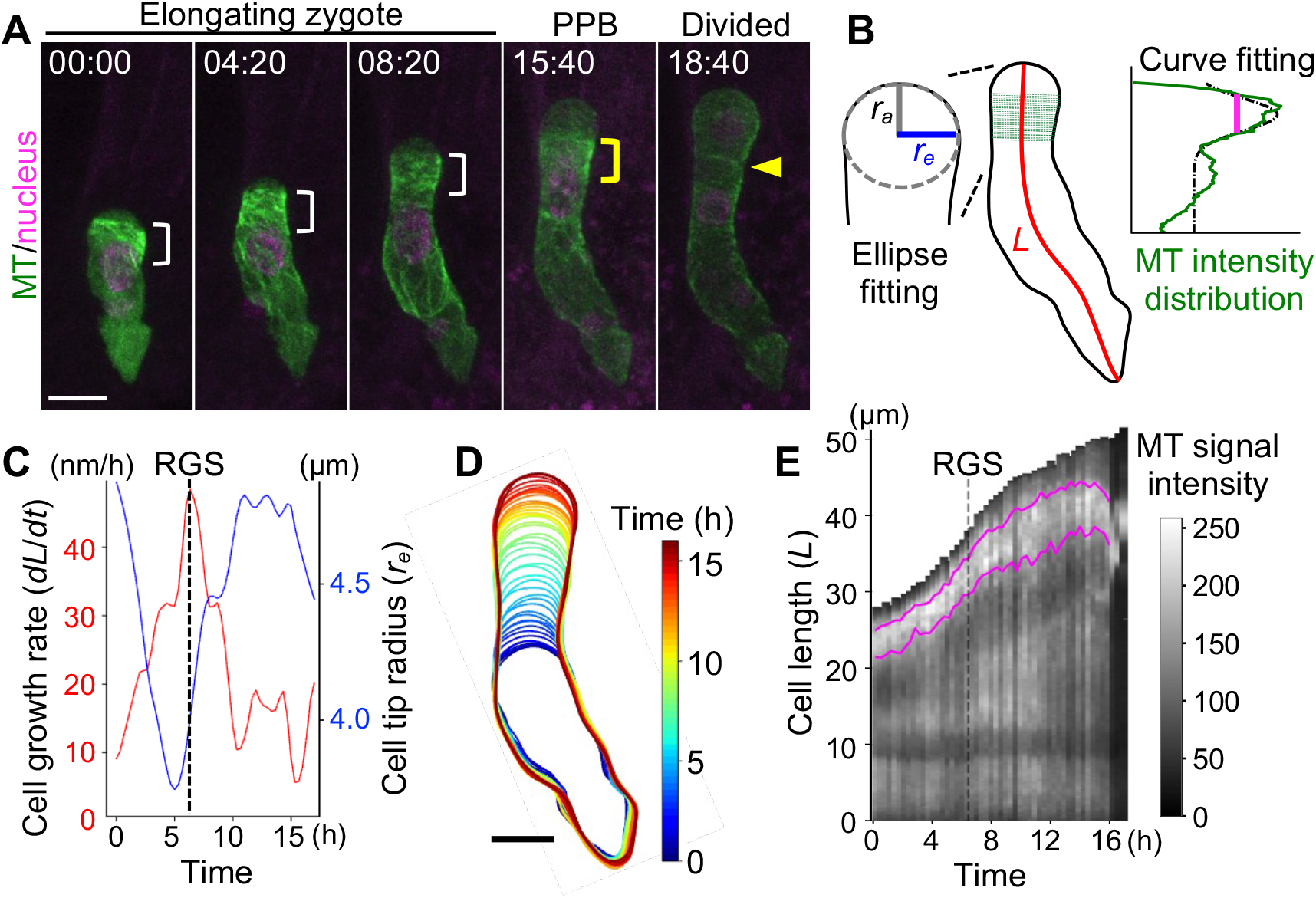
MT bands are kept in a constant position from the growing zygote tip. (**A**) Time-lapse images of a zygote expressing MT (green) and nuclear (magenta) markers. Maximum-intensity projection (MIP) images of two-photon excitation microscopy (2PEM) images are shown. Numbers indicate the time (hour:min) from the first frame. White and yellow brackets show the subapical MT band and PPB, respectively. Yellow arrowhead indicates the cell division plane. (**B**) Schematic illustration of the method used for image quantification. For details, see the Materials and Methods. (**C**) Time course of cell growth rate (*dL/dt*; red) and tip radius (*r*_*e*_; blue) of the zygote shown in A. The rapid growth stage (RGS) is indicated by a dashed line. (**D**) Contour dynamics of the zygote with the time indicated by color code. (**E**) Kymograph showing MT signal intensity projected onto the zygote centerline. Magenta lines indicate the upper and lower ends of the MT band, as determined by Gaussian fitting (as shown in B and Movie S1). Scale bars: 10 μm.

### The subapical MT band acts as a mechanical frame to maintain the shape of the zygote tip

Cortical MTs guide the cellulose synthase complex during cell wall production (*9*). The egg cell lacks an obvious cell wall in the apical region, where the plasma membrane fuses with the sperm cell membrane (*10*). To explore the relationship between the subapical MT band and cell wall production, we used an MT marker and the cellulose-specific dye Calcofluor White (CFW) to observe MTs and cellulose in the same zygote (Fig. 2A and Fig. S3). There was a distinct cellulose signal in the MT band region and below, whereas no clear staining was observed above the band.

**Figure 2.**
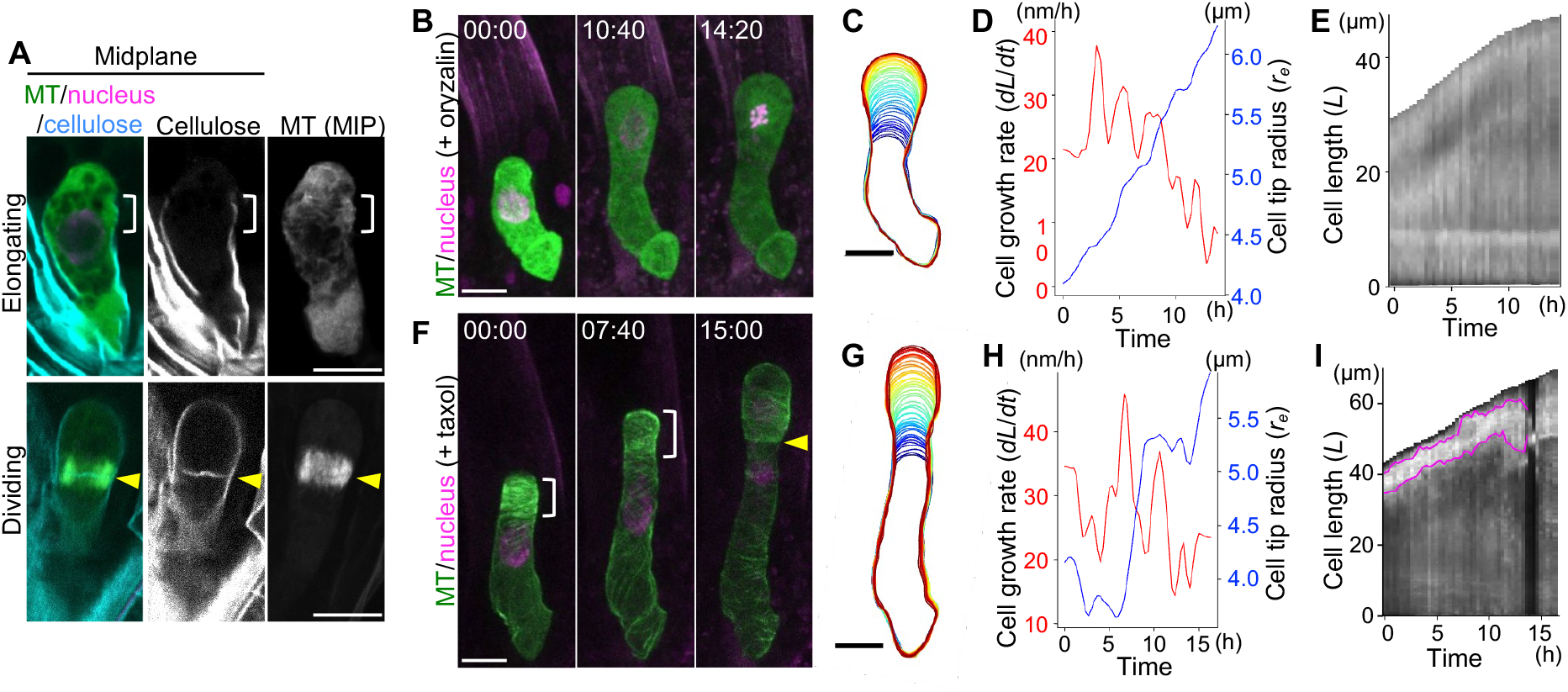
The subapical MT band acts as a mechanical frame to maintain the shape of the zygote tip. (**A**) 2PEM images of cleared zygotes expressing MT (green)/nuclear (magenta) markers at the indicated stages. Cellulose was stained by CFW (cyan). Left and center panels show midplane images, and right panels show MIP images. Brightness was adjusted in the single-color images. Brackets show the MT band position, and yellow arrowheads indicate the new cell plate with phragmoplast. (**B**-**I**) Effects of treatment with a MT polymerization inhibitor (1 μM oryzalin) (B-E) and depolymerization inhibitor (10 μM taxol) (F-I). (B and F) Time-lapse 2PEM images. Contour dynamics (C and G), time course of cell growth rate (*dL/dt*; red) and tip radius (*r*_*e*_; blue) (D and H), and kymographs showing MT signal intensity (E and I) of the zygote shown in B and F, respectively. No MT band was detected in oryzalin-treated zygotes due to diffused signals, as shown in Movie S2. Magenta lines in I indicate the upper and lower ends of the MT band; its precise area could not be extracted by Gaussian fitting due to the distorted shape of the signal peak, as shown in Movie S3. Scale bars: 10 μm.

In dividing zygotes after the MT band disappeared, a clear cellulose signal was observed throughout the cell outline, including the cell apex and newly forming cell plate. These results raise the possibility that MT band–dependent cellulose synthesis forms a rigid cell wall on the lateral side of the zygote, which prevents the radial cell expansion, resulting in the polar zygote elongation.

To test this idea, we analyzed the effects of a MT polymerization inhibitor (oryzalin) and a MT depolymerization inhibitor (taxol). The oryzalin-treated zygotes swelled, with a continuous increase in cell radius and a decrease in growth rate, revealing the loss of radial growth restriction (Fig. 2B-D, Fig. S4, and Movie S2) (*5*). MT bands disappeared and were not detectable by Gaussian fitting (Fig. 2E, Fig. S4C and F, and Movie S2). By contrast, taxol treatment distorted the cell shape, leading to non-uniform growth (Fig. 2F-H, Fig. S5A-F, and Movie S3). The MT bands occasionally became wider and reached toward the cell apex (Fig. 2I, Fig. S5C and F, and Movie S3). The RGS was not detected in either oryzalin- or taxol-treated zygotes, confirming the failure of the spatiotemporal regulation of cell growth. Cell division failed in oryzalin-treated zygotes (Fig. 2B), likely due to blocked formation of the mitotic apparatus. Taxol led a more symmetric than usual cell division, producing a longer apical cell (Fig. 2F and Fig. S5G), implying that the MT band helps determine asymmetric cell division.

In summary, our results indicate that the MT band is maintained at the subapical region of the zygote by the proper balance of polymerization and depolymerization and acts as a mechanical frame to restrict the zygote growth direction, possibly by guiding cell wall synthesis.

### The MT band is located at the subapical region where the material derivative of surface tension reaches its maximum

Next, we explored how the MT band is maintained at a constant position in the actively renewing zygote tip. MTs are proposed to act as a tension sensors (Hoermayer et al. 2024; Moulia et al. 2021), and thus, cortical MT alignment can change in response to cell surface tension. Therefore, we estimated the mechanical properties of growing zygotes by combining computer modeling with time-lapse imaging. We recently found that zygote growth can be reconstructed using the viscoelasto-plastic deformation model (*11*), which simulates cell morphogenesis (deformation from the initial to next shape) based on two mechanical parameters: the expansion force from inside the cell (turgor pressure: *P*); and the deformability of the cell surface (surface extensibility: *Φ*) (Fig. 3A and Fig. S6, see Materials and Methods) (*12*). Although surface extensibility is a complex variable with spatial distribution, we determined that a single parameter defining the range of extensible area (extensibility range: *l*_*g*_), combined with turgor pressure, is sufficient to describe hemispherical tip deformation of the zygote (Fig. S6C and D) (*11*). Therefore, we searched for a suitable combination of *P* and *l*_*g*_ at each time point that explains the actual dynamics of zygote deformation, i.e., changes in the growth rate and tip radius obtained from the time-lapse data, and reconstructed the model zygote (compare ‘Data’ and ‘Model’ in Fig. 3B, and Fig. S7A-C). Based on the model zygote, we calculated the spatial distribution of circumferential surface tension (*σ*_*θ*_) with respect to the zygote apex. The value below the subapical region was clearly larger than that at the tip at all time points examined (Fig. 3C and Movie S4). To precisely measure the temporal change in surface tension, we calculated the material derivative (*Dσ*_*θ*_*/Dt*), considering the zygote tip as a moving coordinate system of surface points (see Materials and Methods). The material derivative markedly increased at the subapical region, which corresponds to the MT band position (Fig. 3D and Movie S4). These features were observed in all tested samples (Fig. S7D-Q), leading to the hypothesis that the MT band is maintained at the subapical region by the large constant material derivative of surface tension. Moreover, the temporal average of MT band width (6.6 µm, Fig. 3E) was wider than the region of maximum material derivative (2.2 µm, Fig. 3D). This observation is consistent with our agent-based model simulations of cortical MT organization (Materials and Methods) where MT bands can form that are wider than the external cue zone mimicking the large material derivative region (Fig. S8).

**Figure 3.**
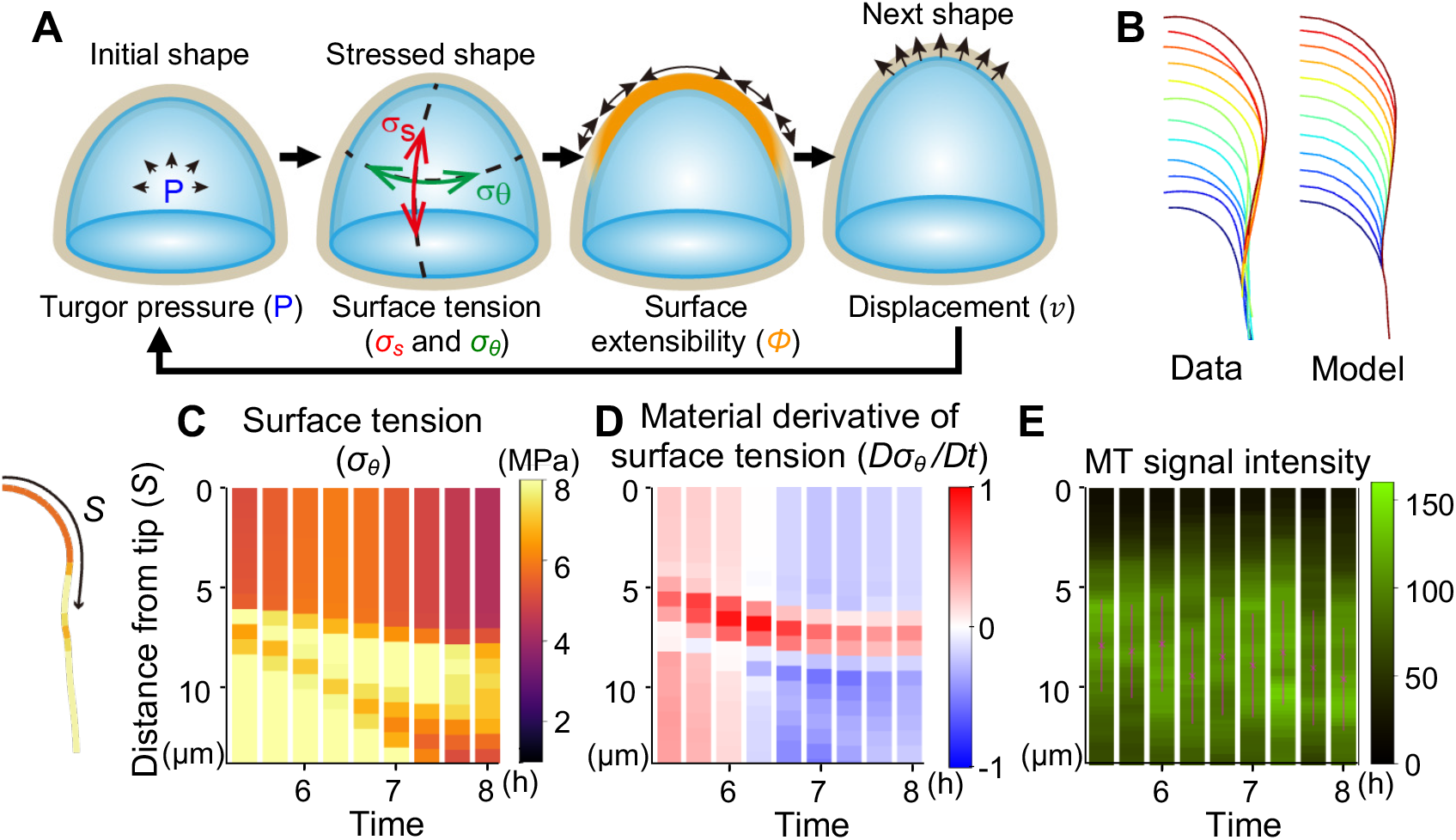
MT bands are present at the subapical region where the material derivative of surface tension reaches its maximum. (**A**) Schematic illustrations of the viscoelasto-plastic deformation model. For additional details, see Materials and Methods. (**B**) Contour dynamics obtained by time-lapse observation of the zygote shown in Fig. 1A (Data) and the reconstructed model zygote (Model). Only the right half of the zygote tip is shown. (**C**-**E**) Calculated circumferential surface tension (*σ*_*θ*_) (C), its material derivative (*Dσ*_*θ*_*/Dt*) (D), and observed MT signal intensity (E), as shown in Movie S4. Values at each distance from the zygote tip (*S*) are plotted with color codes, as shown to the right of each graph. Magenta lines and crosses in E indicate the MT bands and their centers, respectively.

### Continuous zygote elongation is required to maintain the subapical MT band

To experimentally test the importance of growth-dependent changes in tension for MT band maintenance, we used an Arabidopsis mutant in the mitogen-activated protein kinase kinase kinase YODA (YDA), *yda-2991*, in which zygote growth arrests prematurely (*13, 14*). Live-cell imaging of *yda* showed that the subapical MT band was present in young zygotes (Fig. 4A and B, and Movie S5). However, after cell elongation slowed, MT signals dispersed, and thus the MT band became undetectable (visually and by Gaussian fitting). We calculated the surface tension of *yda* zygotes by reconstructing a model zygote and found that MT bands were maintained at the subapical region as long as cell elongation caused a large material derivative of tension; once this large material derivative disappeared, the MT band also became undetectable (Fig. 4D-F, Fig. S9A-C, and Movie S6). These features were observed in all tested samples, and the MT band disappeared within a few hours (∼1–4 hours) after the loss of the large material derivative (Fig. 4D-F, Fig. S9D-S). These results support the importance of cell growth-driven changes in tension for the maintenance of the subapical MT band.

**Figure 4.**
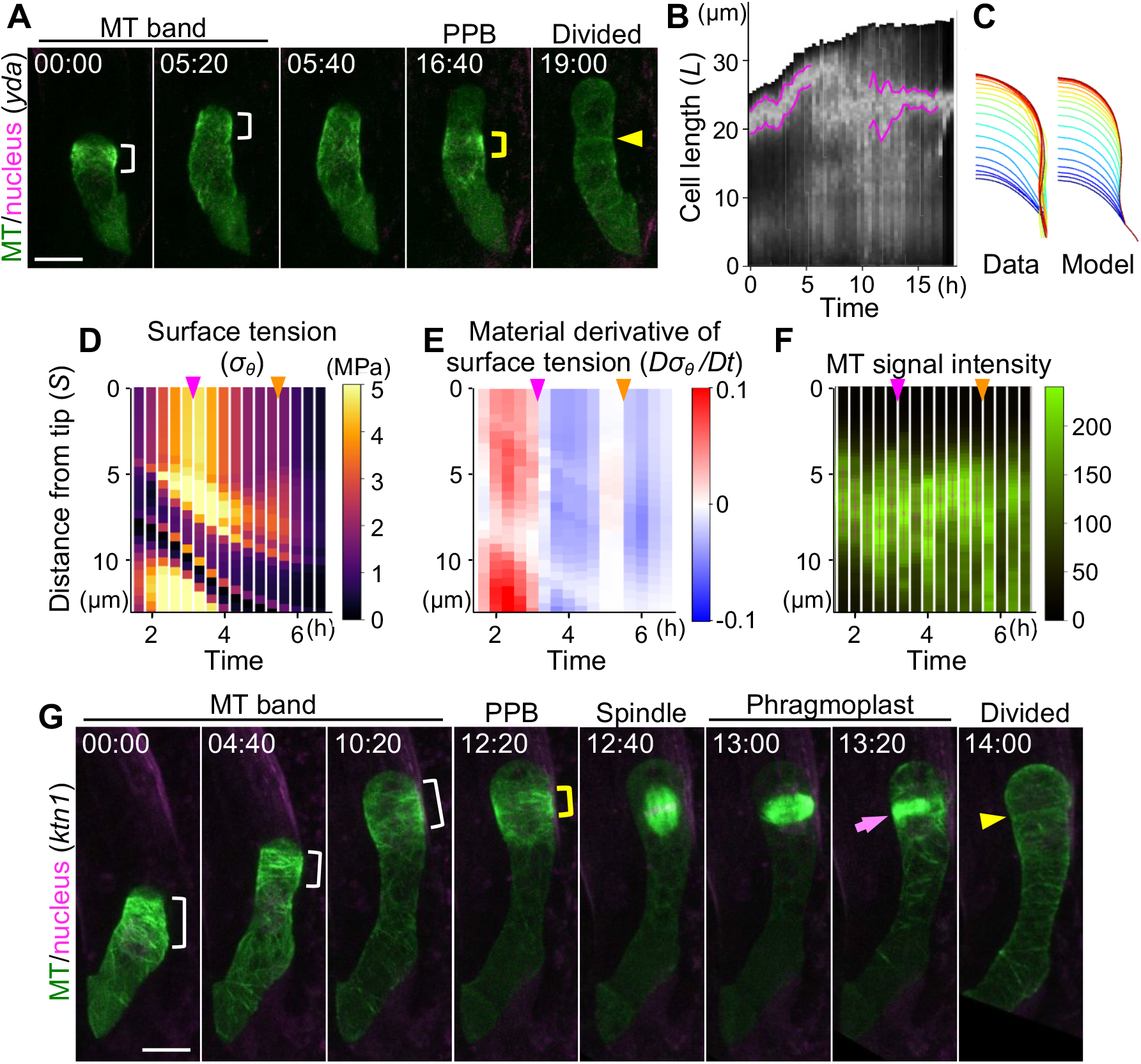
The MT band disappears after zygote elongation ceases. (**A**) Time-lapse images of the *yda* zygote expressing MT (green)/nuclear (magenta) markers. White and yellow brackets show the MT band and PPB, respectively. Yellow arrowhead indicates the cell division plane. (**B**) Kymograph showing MT signal intensity. Magenta lines indicate the MT band based on Gaussian fitting, showing the disappearance of the subapical MT band after 5:20 and the appearance of the PPB at 10:40, as shown in Movie S5. (**C**) Contour dynamics obtained from the time-lapse images shown in A (Data) and the reconstructed model zygote (Model). (**D**-**F**) Circumferential surface tension (*σ*_*θ*_) (D), its material derivative (*Dσ*_*θ*_*/Dt*) (E), and MT signal intensity (F), as shown in Movie S6. Magenta lines and crosses in F indicate the MT bands and their centers, respectively. Magenta and orange arrowheads indicate when the large material derivative of surface tension is lost and when the MT bands are no longer detectable, respectively. (**G**) Time-lapse images of the *ktn1* zygote expressing MT (green)/nuclear (magenta) markers, as shown in Movie S7. Magenta arrow indicates the delayed phragmoplast. Scale bars: 10 μm.

Although it has been proposed that MTs can autonomously sense tension via tubulin itself, some regulatory factors are also known to rearrange MT patterns in response to tension; a well-known factor is the MT-severing enzyme KATANIN1 (KTN1) (*15, 16*). Therefore, we examined the behavior of zygotes of the KTN1 mutant *ktn1-2*, which shows abnormal cell division patterns during embryogenesis (*17*), using live-cell imaging. None of the zygotes produced in *ktn1* heterozygous mother plants exhibited detectably abnormal MT band maintenance, cell elongation, or subsequent PPB formation (0%, n=20), even in zygotes that showed delayed phragmoplast expansion, as reported as the *ktn1* phenotype in other tissues (Fig. 4G and Movie S7) (*18*). These results indicate that KTN1 is not required for the mechanical feedback of cell growth and MT organization.

### MT bands help determine the cell division site in cooperation with the nucleus

In *yda* mutant zygotes, the PPB appeared after the disappearance of the subapical MT band (Fig. 4B, Fig. S9D and L, and Movie S5). The PPB formed away from the subapical region, resulting in a more symmetric than usual cell division. Since MT stabilization due to taxol treatment also affected asymmetric cell division (Fig. 2F and Fig. S5G), we tested whether the MT band is involved in determining the position of the PPB by examining the positional relationships among the MT band, PPB, nucleus, and cell division plane in wild-type zygotes (Fig. 5A and B). Before cell division, the PPB appeared slightly below the region containing the subapical MT band, above the nucleus (Figs. 1E and 5A, and Movie S1). The gap was subtle but consistent, as shown by significant differences of the center positions of the MT band, PPB, and nucleus (Fig. 5B). The spindle formed between the PPB and nucleus, establishing the cell division plane (Fig. 5A). Indeed, the cell division site was significantly different from the sites of all these structures, except for the spindle (Fig. 5B), suggesting that the division site is determined by spindle formation between the nucleus and PPB. To investigate whether the two transverse MT structures, the subapical MT band and the PPB, are generated by a common molecular mechanism, we focused on the core regulator of PPB formation, TON1 RECRUITING MOTIF (TRM), whose triple mutant (*trm678*) lacks a visible PPB (*19*). The subapical MT band was retained by *trm678* zygotes during cell elongation, but dispersed without forming the PPB before cell division (Fig. 5C and Movie S8). The spindle and phragmoplast formed at the nuclear position, and their shapes were unstable, highlighting the importance of the PPB for supporting these structures. A cell division plane formed at the same site, resulting in an apical cell that was a bit longer than the wild type (Fig. 5D). These results indicate that MT bands and PPBs form via different molecular mechanisms. This notion is consistent with the finding that in *yda* zygotes, the PPB formed in a different position several hours after the MT band disappeared (Fig. 4A and B). Taken together, our data indicate that PPB formation near the MT band site is important to ensure accurate asymmetric zygote division in wild-type Arabidopsis.

**Figure 5.**
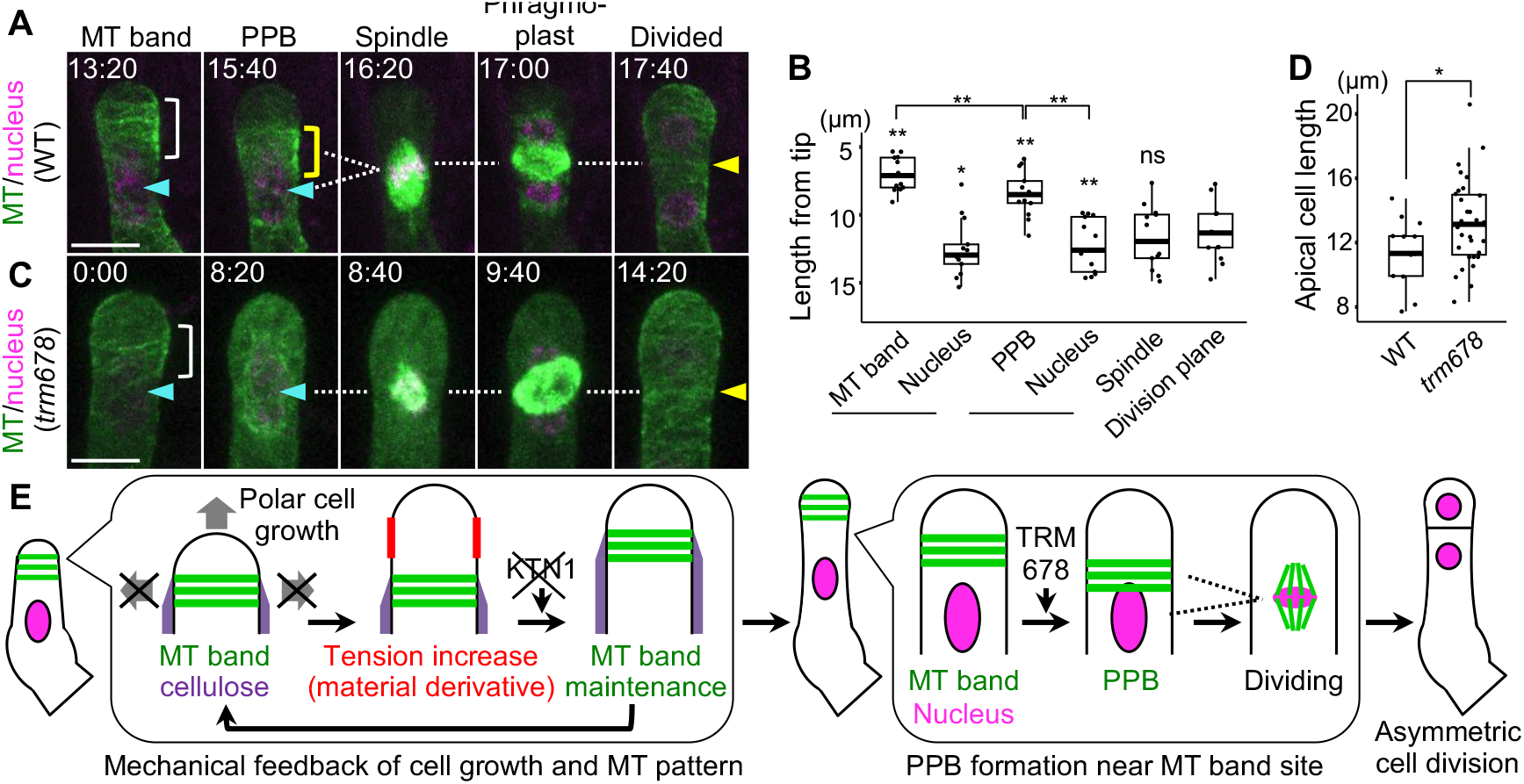
The division plane forms between the MT band and nucleus. (**A**) Time-lapse images of a wild type (WT) zygote expressing MT (green)/nuclear (magenta) markers. Enlarged images of the zygote in Fig. 1A are shown. The PPB (yellow rectangle) forms below the MT band (white rectangle) and above the nucleus (cyan arrowhead). The cell division plane then forms between the PPB and nucleus (yellow arrowhead and dashed lines). (**B**) Boxplot showing the center positions of the indicated structures. The locations of the MT band and PPB are shown together with the location of the nucleus, as measured at the same time points. Significant differences in the location of each structure relative to the cell division plane, as well as the MT band, PPB, and nucleus, were determined by Wilcoxon’s signed rank test (n=13). n.s., not significant; \**P* < 0.05 and \*\**P* < 0.01. (**C**) Time-lapse images of the *trm678* zygote. (**D**) Boxplot of apical cell lengths of WT and *trm678* zygotes. Significant differences were determined by single-sided Welch’s T-test (n=13 for WT, n=31 for *trm678*). \**P* < 0.05. The WT zygotes correspond to the “division plane” samples in Fig. 5B. (**E**) Schematic representation of dynamics and the proposed mechanism. Scale bars: 10 μm.

## Discussion

We propose an autonomous feedback mechanism whereby polar zygote growth produces a large material derivative of surface tension, which in turn restricts the zygote growth direction via MT band formation, ultimately leading to accurate asymmetric cell division (Fig. 5E).

In most animal and plant cells, the dynamic behavior of the cortical cytoskeleton is crucial for growth dynamics. In plants, the rigid cell wall generally covers the entire cell surface. However, we determined that the zygote tip lacks cellulose, which is in agreement with the fact that the sperm cell membrane fuses to the top of the egg cell (*10, 20*). The MT band maintains the boundary between the apical region (lacking cellulose) and the basal region (containing cellulose), and acts as a mechanical frame to restrict the direction of zygote growth (Fig. 5E). Since the membrane fusion of the sperm and the egg is not specific to Arabidopsis, and zygotes are elongated and undergo transverse division in various plant species, such as the angiosperm tobacco (*Nicotiana tabacum*) and the brown alga *Dictyota* (*21, 22*), it would be important to determine whether MT band-dependent zygote growth occurs in other species.

We also showed that cell growth-dependent MT alignment occurs via surface tension. The MTs were proposed to function as a tension sensor in various experiments. For example, in an *in vivo* experiment, forcing protoplasts to change shape led to MT rearrangement (*23*). In the protoplasts, the tension-induced MT pattern was reoriented after several hours, indicating that MTs can respond to temporal changes in tension, rather than the absolute magnitude of tension, with a time lag. Similarly, in the current study, the MT band in *yda* zygotes disappeared a few hours after the loss of a large material derivative of surface tension. In plant tissues, where neighboring cells are tightly anchored by cell walls, damage to and growth of adjacent cells cause external changes in tension and promote MT-dependent cell morphogenesis to adjust the overall tissue shape (*24, 25*). By contrast, the zygote is surrounded by a fluidic giant cell, the endosperm, and thus would freely grow without significant external forces. Therefore, the zygote may induce internal changes in tension by regulating its own growth rate and local cellulose synthesis, resulting in cell polarization and thus asymmetric cell division. In such stand-alone situation, it seems reasonable that the zygote adopts a simple mechanical mechanism to autonomously amplify subtle polarity information, such as sperm entry from the apical end, leading to the formation of apical–basal axis. This notion of concise autonomy is consistent with the finding that zygote growth does not require the MT regulator KTN1, which acts in response to exogenous stimuli such as external mechanical stress and blue light irradiation as well as developmental regulations involving hormonal signaling (*16, 26, 27*). Additional studies are needed to examine whether the MT itself responds to tension, or whether mechanosensitive sensors, such as the calcium ion channel MCA2, which is activated by membrane tension, are involved (*28*).

We also showed that the MT band can influence PPB position and thus contribute to the asymmetry of cell division (Fig. 5E). This seems odd, since the PPB usually forms around the nucleus, but pre-adjusting PPB position via MT patterning is an effective strategy to ensure asymmetric division. Indeed, in a recent study, polarly localizing proteins in the leaf epidermis deplete stable MTs from a particular cell domain and thus limit the area in which the PPB could form (*29*). Nevertheless, this epidermis cell divides at the nuclear position (*30*), highlighting the uniqueness of the zygote that forms the PPB and the division plane at a different location than the nucleus. We can only speculate about the mechanisms that determine the exact position of the PPB in zygotes. As mentioned above, the pre-depletion of MTs inhibits PPB formation (*29*). Therefore, it appears to make sense that the PPB forms near the MT band site, an abundant source of MTs. The *trm678* zygotes divided at the nuclear position, forming longer apical cells than the wild type. Since apical cell geometry is crucial for setting proper transverse division, an early event in radial axis formation (*31*), the MT band-dependent setting of the division site in the zygote would be important for the formation of a small spherical apical cell for subsequent embryo patterning.

Our findings highlight the importance of mechanics, especially surface tension, in the dynamics of zygote development, representing the initial event of plant ontogeny. Therefore, our findings will facilitate the interdisciplinary researches of plant biology with mechanical engineering, physics, and data science, which would help the manipulation of seed development to enhance agriculture.

## Materials and Methods

### Plant growth conditions and transgenic lines

All Arabidopsis lines are in the Col-0 background. The *yda-2991, ktn1-2* (SAIL_343_D12), and *trm678* triple mutants were described previously (*14, 19, 32*). The MT marker (EC1p::Clover-TUA6: coded as MU2228) was same as previously described (*5*), only except that it is in pMDC99 binary vector (*33*). In MT/nuclear marker, EC1p::Clover-TUA6 was combined with a nuclear marker ABA INSENSITIVE4 (ABI4)p::H2B-tdTomato (coded as MU2463), which contains the 981-bp ABI4 promoter (*34*), the full-length coding region of H2B (AT1G07790), tdTomato, and the NOS terminator in the pPZP211 binary vector. Plants were grown at 18–22°C under continuous light or long-day conditions (16-hour light/8-hour dark).

### Time-lapse observations and histological analysis

The zygote live-cell imaging was performed using a laser-scanning inverted microscope (A1; Nikon) equipped with a pulse laser (Mai Tai DeepSee; Spectra-Physics) and (AX; Nikon) equipped with a pulse laser (InSight X3 Dual option; Spectra-Physics) as previously described (*35, 36*). Time-lapse images were acquired every 20 min at 25 or 31 *z*-stacks with 1-μm intervals and using a 40× water-immersion objective lens (CFI Apo LWD WI; Nikon) with Immersol W 2010 (Zeiss) immersion medium. Fluorescence signals were detected by the external non-descanned GaAsP PMT detectors. In A1 system, we used two dichroic mirrors (DM495 and DM560), a short-pass filter (492 nm/SP) for CFW stain (see below), a band-pass filter (525/50 nm) for Clover, and a mirror for tdTomato. In AX system, we used three dichroic mirrors (DM488, DM560 and DM685), and three band-pass filters (457/70 nm for CFW stain, 525/50 nm for Clover, and 605/70 nm for tdTomato). For inhibitor treatment, 0.1% DMSO and individual inhibitors [0.1% DMSO containing 1 μM oryzalin (Chem service) or 10 μM taxol (Wako, Paclitaxel)] were added to the media ∼1 h before observation, as previously described (*37*).

For cell wall staining, tissue clearing was performed using ClearSeeAlpha solution (*38*). The self-pollinated pistils were fixed with 4% (w/v) PFA for 1 hour in PBS under vacuum (∼690 mmHg) at room temperature. Fixed pistils were washed in PBS and cleared with ClearSeeAlpha for 3-5 days. Then the cleared pistils were stained with 1 mg/ml CFW (Wako) in water overnight and then mounted in ClearSeeAlpha droplet. The observation was performed with the above microscope (AX). The images were acquired at 25 z-stacks of 1-μm intervals using the above objective lens. Maximum intensity projection (MIP) and image processing were performed using NIS-Elements software (Nikon), or Fiji (https://fiji.sc/).

### Quantification of growth rate and tip shape from live-cell imaging sequence

The location of the zygote in ovules can be fluctuated due to floating and ovule growth in liquid cultivation medium. Based on the previously developed coordinate normalization (image-based and contour-based) methods to fix the zygote position (*2*), we developed method, i.e., the improved Contour-based Coordinate Normalization (improved CCN), using the normalized cross-correlation function(*39*) between the image at the reference time and that at time t using a parallel shift (*u, v*) and a rotation *θ* as follows.

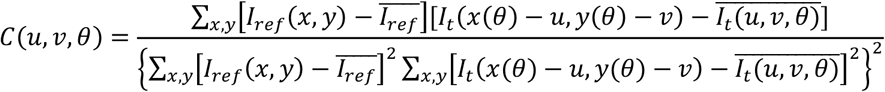

where *I*_*ref*_(*x, y)* is the fluorescence intensity of the reference image (typically at the initial time frame) at pixel x and y, 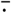 is the spatial average of ⋅, and *I*_*t*_ is the modified image at time t. We applied an exhaustive search for the optimal set (*u*_*opt*_, *v*_*opt*_, *θ*_*opt*_) which minimizes the normalized cross-correlation function.

Using the improved CCN, we first extracted cell contours from the raw data for all the image sequences using binarization in Fiji (Fig. S1A and S1B). Next, we applied the Voronoi diagram (Fig. S1C) which evaluates the skeleton inside the contour as previously analyzed (*40*). As this method may create lateral branches, we defined the part of the centerline as the longest skeleton neglecting all the branches shorter than the longest one (illustrated with the red line in Fig. S1D). We then smoothly extended the ends to the contour (the dashed green line in Fig. S1D). We quantified the growth rate *dL/dt* and tip shape (*r*_*e*_, *r*_*a*_) by measuring the cell length *L* and the ellipse fitting with the horizontal radius *r*_*e*_ and the vertical radius *r*_*a*_ (Fig. S1E).

### Detection of MT band using truncated Gaussian fitting

Based on the cell shape and centerline described above (Fig. S1F), we projected the fluorescence intensity of MT marker within the cell to the centerline (Fig. S1G). We then plotted the MT signal intensity as a function of the position z along the centerline (Fig. S1H). We fitted the intensity with the truncated Gaussian distribution using parameters (*α, β, μ, σ, b*) with average *μ* and standard deviation *σ* of the distribution as follows.

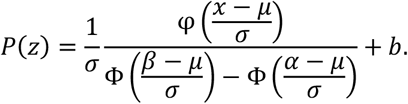

Here, the Gaussian probability density function is 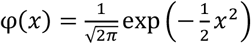 and the cumulative distribution function is 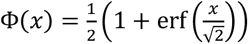. We then defined the MT band by *σ* from the average, 2*σ* in width. Using the extracted *μ* and *σ*, we could plot the line segment of the MT band as shown as the magenta line in Fig. S1I. The coefficient of determination (*r*^2^) is calculated as follows.

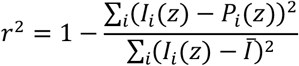

where *I*_*i*_(z) stands for the i-th fluorescent intensity at the position z along the cell centerline greater than *b*, and 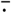 is the spatial average of ⋅.

### Construction of model zygote based on viscoelastic-plastic deformation model

We constructed a mechanical model for a zygote elongation based on the model previously developed for root hairs as schematically illustrated in Fig. S6A (*11, 12*). This model describes the degree of surface stress derived from the pressure on the input shape (curvature κ and cell wall thickness δ) as shown in the arrow from shape to stress. Then, it calculates the strain rate 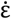 which is the consequence of the wall yield beyond the yield stress *σ*_y_ (the arrow from stress to strain). Finally, the next shape is achieved by applying the displacement vector *v* on the current shape (the arrow from strain to shape).

Cell surface mechanical stresses in the meridional and circumferential directions were formulated as,

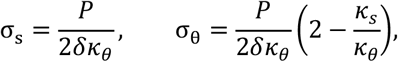

respectively. Here, the circumferential curvature *κ*_*θ*_ was calculated by 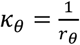 where the radius *r*_*θ*_ is the curvature of the surface lying on the plane perpendicular to the meridional coordinate (Fig.S6B). On the other hand, the meridional curvature *κ*_*S*_ was calculated by 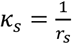 where the radius *r*_*S*_ is the curvature perpendicular to the meridional coordinate *S* (Fig. S6C).

The strain rates 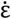 on the surface in the meridional and circumferential directions were formulated as,

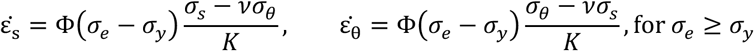

respectively, where Φ(S) is the cell surface extensibility defined as 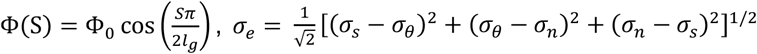 with *σ*_*n*_ standing for the stress perpendicular to the surface, 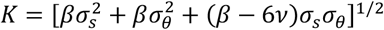 and 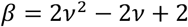. Here, *l*_*g*_ is the range of extensible area (extensibility range), and Φ_0_ is the magnitude of extensibility at *S* = 0. We note that the strain rate becomes zero for *σ*_*e*_ < *σ*_*y*_, and *σ*_*n*_ makes the cell wall thinning due to stretching with *v* ≤ 1 but we assumed that new wall material maintains wall thickness in this study. Since we found that the zygote tips are hemispherical (*2*), we utilized cosine function to describe surface extensibility based on the knowledge in case of hemisphere-like shape (*41, 42*). The extensibility means the degree of deformability as shown in Fig. S6C. The displacement vectors in the tangential direction and in the perpendicular direction were,

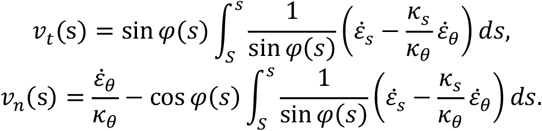

The initial configuration is based on the hemispherical shape with radius larger than the data. The simulation step was smaller than the data time interval 20 min and we solved the above equations per simulation step. Depending on the parameter *P* and *l*_*g*_, the growth speed and cell shape were altered as summarized in Fig. S6D.

### Parameter estimation of turgor pressure *P* and range of extensible area *l*_*g*_

We estimated the parameter set (*P, l*_*g*_) through the quantification of growth and tip radius (Fig. S6E). Plotting analyzed data on the so-called morphospace 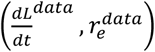 summarizing the characteristic features of the zygote cell shape, we exhaustively searched for the best fitted model parameters (*P, l*_*g*_ ) and resulting 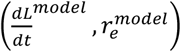 that minimizes the error function 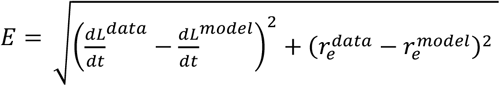. This fitting process was applied to each two sequential time frames.

### Calculation of the material derivative of the circumferential stress

The material derivative of the meridional stress was calculated as,

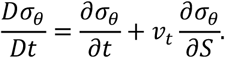

where *v*_*t*_ is the displacement vector in the tangential direction. We note that the time partial derivative is corrected with the tangential displacement depending on the Lagrangian flow of the point on the cell surface.

### Cortical MTs agent-based model

We constructed an agent-based model of the cortical MTs (CMTs) where we took the CMTs as agents (elements) and analyzed the collective behavior of the agents based on the previous works (*43, 44*). We accounted for the following three main dynamics: single MT growth and shrinkage, MTs interaction, and orientational modification based on given large surface tension as an external directional cue.

For single MT growth and shrinkage, we used the growth speed of the plus end v^+ = 0.08 µm/sec and shrinkage speed of the plus end v^-=0.16 µm/sec, and shrinkage speed of the minus end (so-called treadmilling speed) v^tm=0.01 μm/sec. The switching rate from shrinkage to growth at the plus end (so-called rescue rate) is *r*_*r*_ = 0.007 1/sec and the switching rate from growth to shrinkage (so-called spontaneous catastrophe rate) *r*_*c*_ is in the range [0.001-0.01] 1/sec. The nucleation rate of CMTs on the surface *r*_*n*_ = 0.001 1/sec.

For MTs interaction, we considered the interactions between CMTs based on the previous studies (*44, 45*). First, we took account for the zippering effect where the growing CMT changes the direction of the growing front parallel to another CMT with the colliding angle less than 40 degrees. Second, we took account for the induced catastrophe where it begins to depolymerize from the growing front with the probability 50% with the colliding angle more than 40 degrees. Third, we took account for the crossover where the growing CMT crosses another CMT with the probability 50% with the colliding angle more than 40 degrees.

For orientational modification by a directional cue, we defined the directional cue zone (DCZ) where the newly added CMT is biased in a circumferential direction with weight *b*_*d*_.

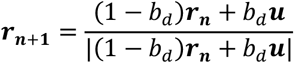

where ***r***_***n***_ and ***r***_***n***+1_ are the current and next unit vector of CMT plus end respectively, ***u*** is a unit vector in the circumferential direction and | ⋅ | is the standard norm of ⋅.

Based on the typical spatial scale of a zygote 5 µm in radius, we set the cylindrical geometry with radius 5 µm. We set the unit of the length of the CMT segment as 8 nm, the width of a ring of tubulin as same as the previous study (*46*). We set the speed of growth at the plus end as 4.8 µm/min (*44*). The simulation time step is 0.2 seconds. Most of the simulation iterates 12000 times corresponding to 40 min in reality. We note that we also implemented the finite tubulin-pool effect as same as the previous study (*43*).

## Acknowledgments

We thank Tamiko Ambo, Hisa Yoshida, Satomi Watanabe, Yuko Kudo, Junko Kato for technical support, Yusuke Kimata, Takumi Higaki and Koichi Fujimoto for helpful discussion, Takema Sasaki for providing *ktn1-2* mutant, and Martine Pastuglia for *trm678* triple mutants. This work was supported by the Japan Society for the Promotion of Science [a Grant-in-Aid for Research Activity Start-up (JP21K20650 to H.M.), a Grant-in-Aid for Early-Career Scientists (JP22K15135 to H.M. and JP20K15832 to S.T.), a Grant-in-Aid for Scientific Research on Innovative Areas (JP19H05670, and JP19H05676 to M.U.; JP16H06280 (Advanced Bioimaging Support)), a Grant-in-Aid for Scientific Research (B) (JP23H02494 to M.U.), and International Leading Research KEPLR (JP22K21352) to M.U.], the Japan Science and Technology Agency [CREST (JPMJCR2121 to S.T. and M.U.)], the Suntory Rising Stars Encouragement Program in Life Sciences (SunRiSE; to M.U.), and the Toray Science Foundation (20-6102 to M.U.).

