## Supplementary material for "Temporal changes in surface tension guide the accurate asymmetric division of Arabidopsis zygotes": Fig. S

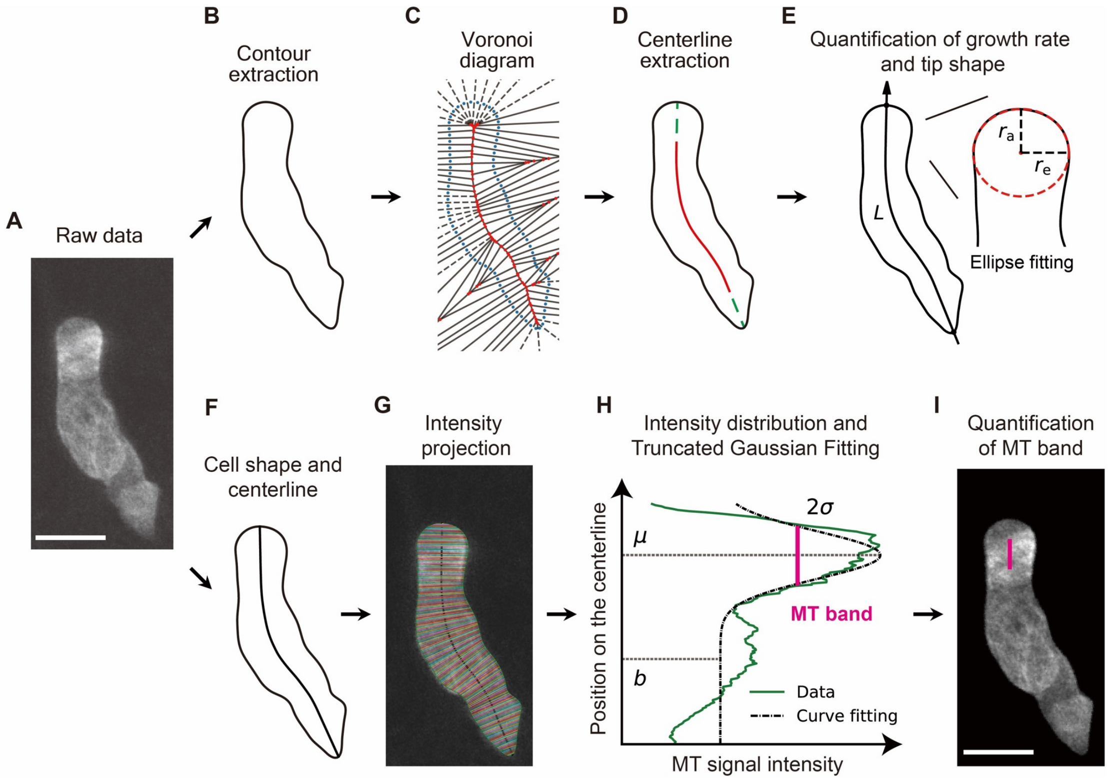


**Fig. S1. Procedure of image processing and quantification analysis.** (**A**-**E**) Schematic illustrations of improved CCN method to quantify the zygote growth dynamics. For additional details, see Materials and Methods. (A) Original 2PEM image of the zygote showing MT marker signal. (B) Contour extraction using binarization in Fiji. (C) Voronoi diagram to evaluate the skeleton patten. (D) Centerline extraction as the longest skeleton in C (red line) and its extension to the cell edge (dashed green line). (E) Quantification of growth rate based on cell length and tip radius measurement by ellipse fitting (dashed red line). (**F**-**I**) Schematic illustrations of truncated Gaussian fitting to detect MT band. (F) Cell contour and centerline as identified above. (G) Intensity projection of MT signal to the centerline in F. (H) Intensity distribution along the centerline (green line) and curve fitting based on truncated Gaussian distribution (dashed black line). MT band region was detected as 2σ (magenta line). (I) Overlay of the extracted band region on the raw image. Scale bars: 10 μm.

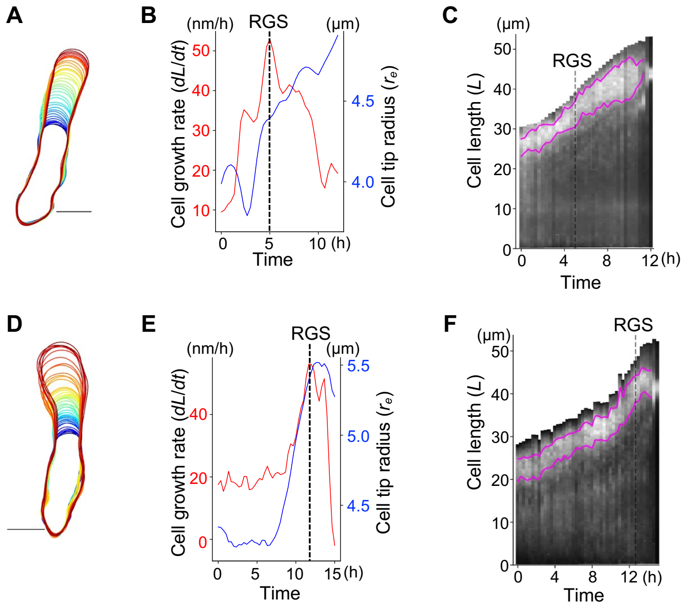


**Fig. S2. Dynamics of MT pattern and cell growth of wild type zygotes.** (**A**-**F**) Two other representative zygotes expressing MT/nuclear marker (A-C and D-F), analyzed as in Fig. 1. Contour dynamics of the zygote (A and D), time course of cell growth rate (*dL/dt*; red) and tip radius ($r_e$; blue) (B and E), and kymograph showing MT signal intensity (C and F). Timing of RGS is shown as dashed lines. Magenta lines indicate the upper and lower ends of the MT band, which is extracted by Gaussian fitting. Scale bars: 10 μm.

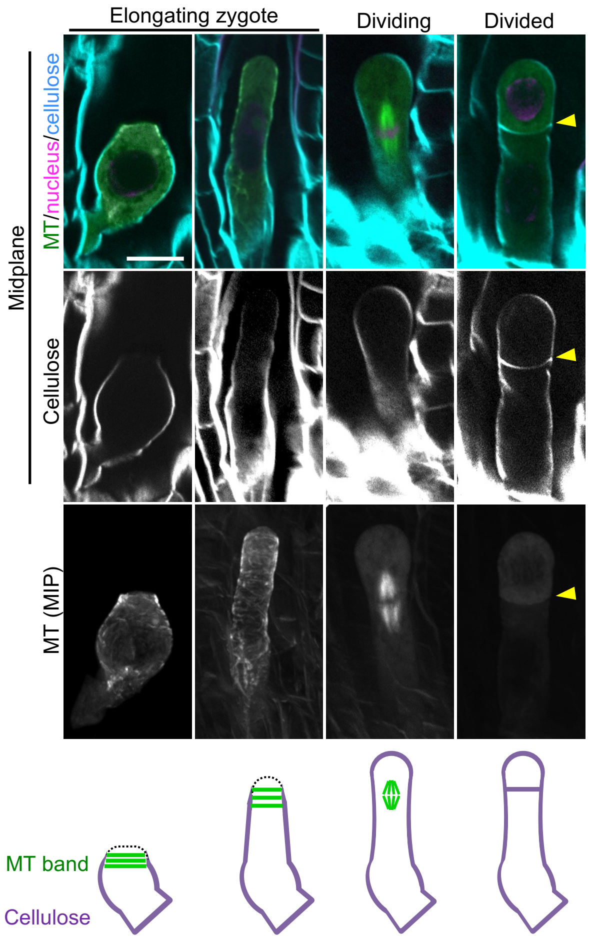


**Fig. S3. Dynamics of cellulose pattern in the zygotes.**
2PEM image of cleared zygotes expressing MT (green)/nuclear (magenta) marker at indicated stages. Cellulose was stained by CFW (cyan). Upper and center panels show midplane images, and lower panels display the MIP images. Dividing zygote contains spindle. Yellow arrowheads indicate the cell division plane. Brightness is adjusted in single-color images. Illustrations show a summary of the respective stages. MT (green) and cellulose (purple) are shown, and the zygote tips with no detectable cellulose signal are indicated by dashed black lines. Scale bar: 10 µm.

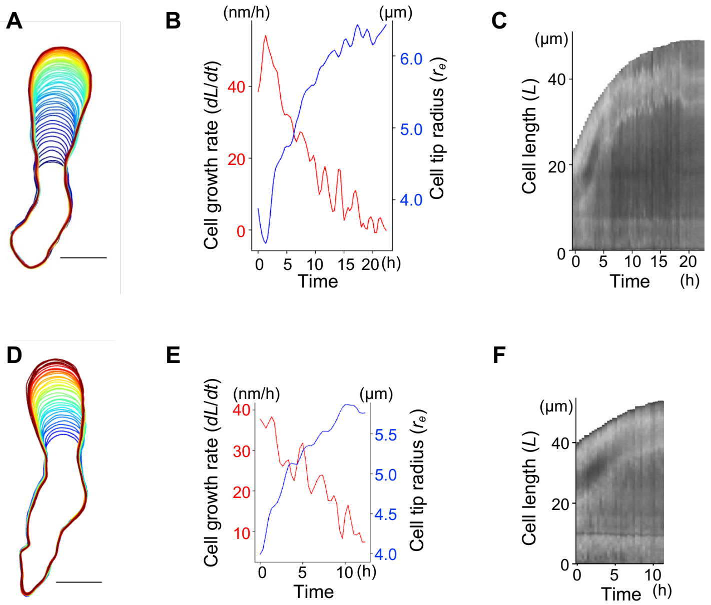


**Fig. S4. Loss of MT pattern and swollen tip in oryzalin-treated zygotes.** (**A**-**F**) Two other representative zygotes expressing MT/nuclear marker in the presence of MT polymerization inhibitor (1 μM oryzalin) (A-C and D-F), analyzed as in Fig. 2. Contour dynamics of the zygote (A and D), time course of cell growth rate (*dL/dt*; red) and tip radius ($r_e$; blue) (B and E), and kymograph showing MT signal intensity (C and F). In both zygotes, RGS, a transient change in growth rate and cell tip radius, was not detected. MT bands were also not detected due to diffuse signal. Scale bars: 10 μm.

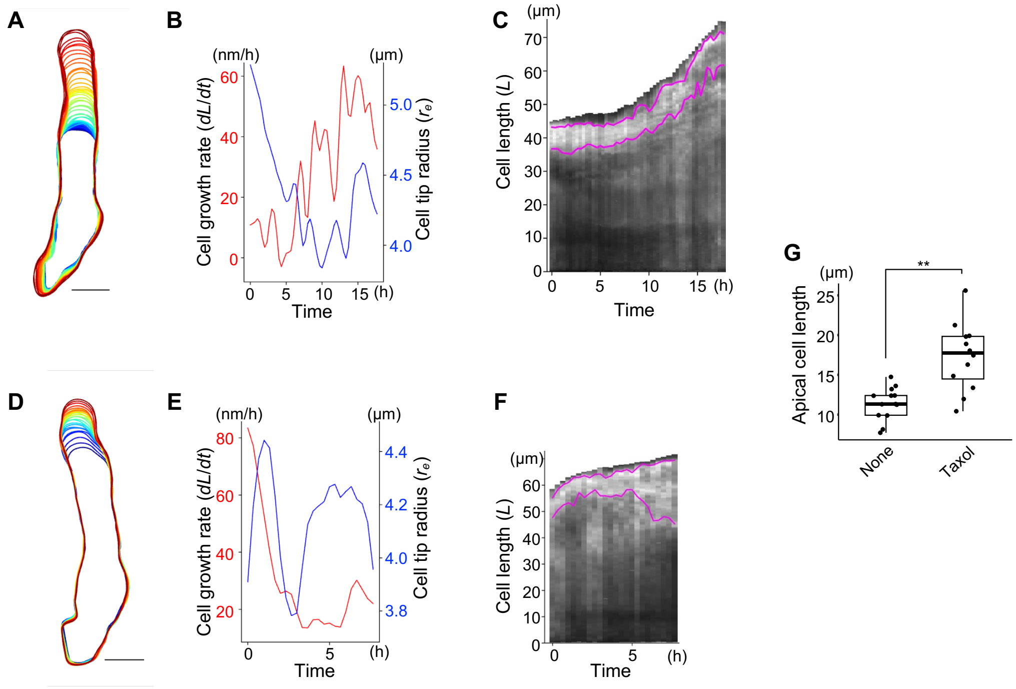


**Fig. S5. Stabilized MT bands and distorted tip in taxol-treated zygotes.** (**A**-**F**) Two other representative zygotes expressing MT/nuclear marker in the presence of MT depolymerization inhibitor (10 µM taxol) (A-C and D-F), analyzed as in Fig. 2. Contour dynamics of the zygote (A and D), time course of cell growth rate (*dL/dt*; red) and tip radius ($r_e$; blue) (B and E), and kymograph showing MT signal intensity (C and F). In both zygotes, RGS, a transient change in growth rate and cell tip radius, was not detected. Magenta lines in C and F indicate the upper and lower ends of the MT bands, but their precise area could not be extracted by Gaussian fitting due to distorted shape of signal peak. (**G**) Boxplot of apical cell lengths of non-treated (none) and taxol-treated (taxol) zygote. Cell length just after the zygote division was measured using time-lapse observation images. Significant differences were determined by single sided Welch's T-test (n=13 for none, n=12 for taxol). ***P* < 0.01. The non-treated zygotes correspond to WT zygotes in Fig. 5B and D. Scale bars: 10 µm.

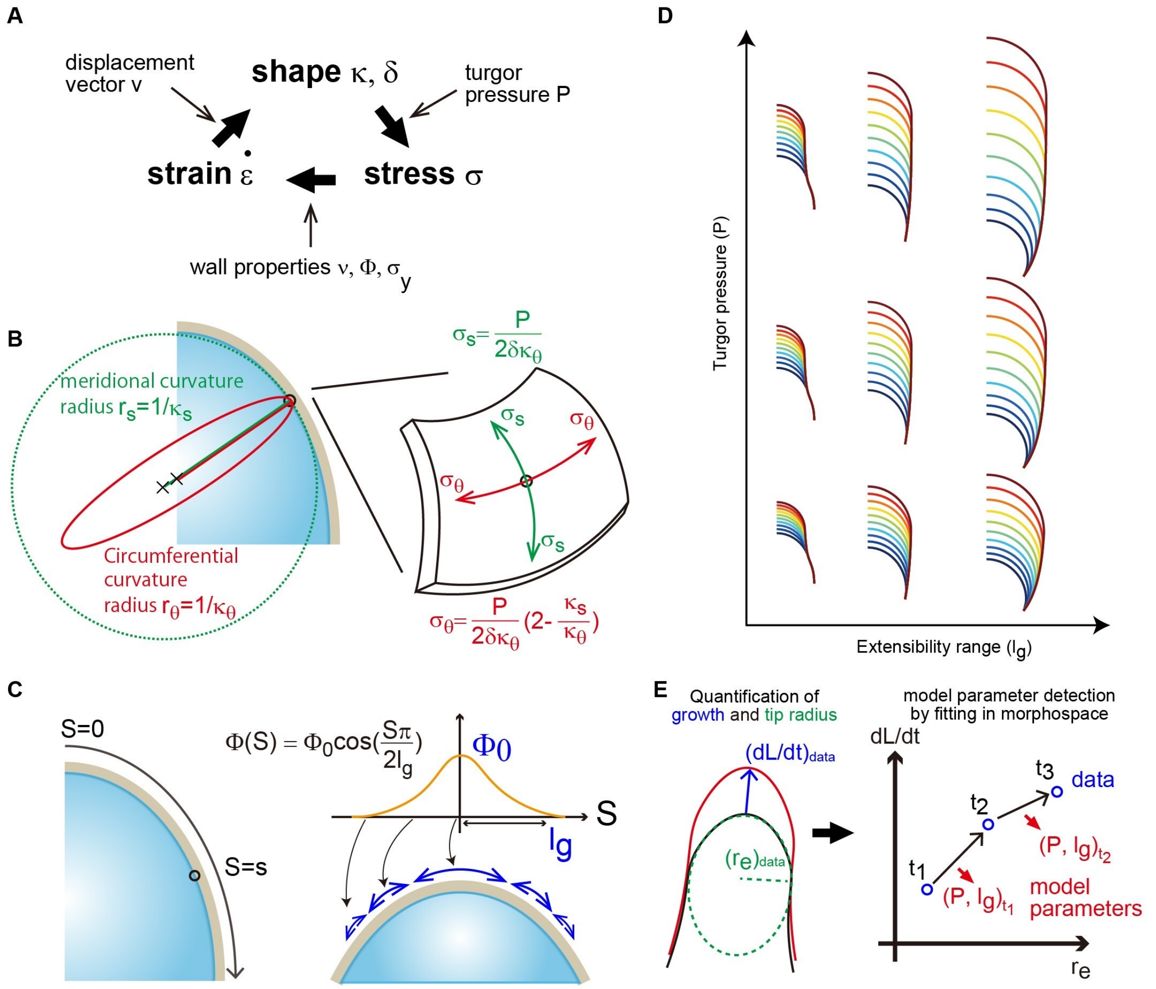


**Fig. S6. Procedure of model zygote construction.** (**A**) Mechanical relations for cell growth in viscoelasto-plastic deformation model. For additional details, see Materials and Methods. (**B**) Cell surface mechanical stress in the meridional direction and in the circumferential direction. (**C**) Schematic illustration of surface extensibility in zygotes with hemispherical tips. (**D**) Variation of simulation results of zygote growth dynamics as functions of $P$ and $l_g$. (**E**) Model parameter detection for each time point based on time-lapse observation data.

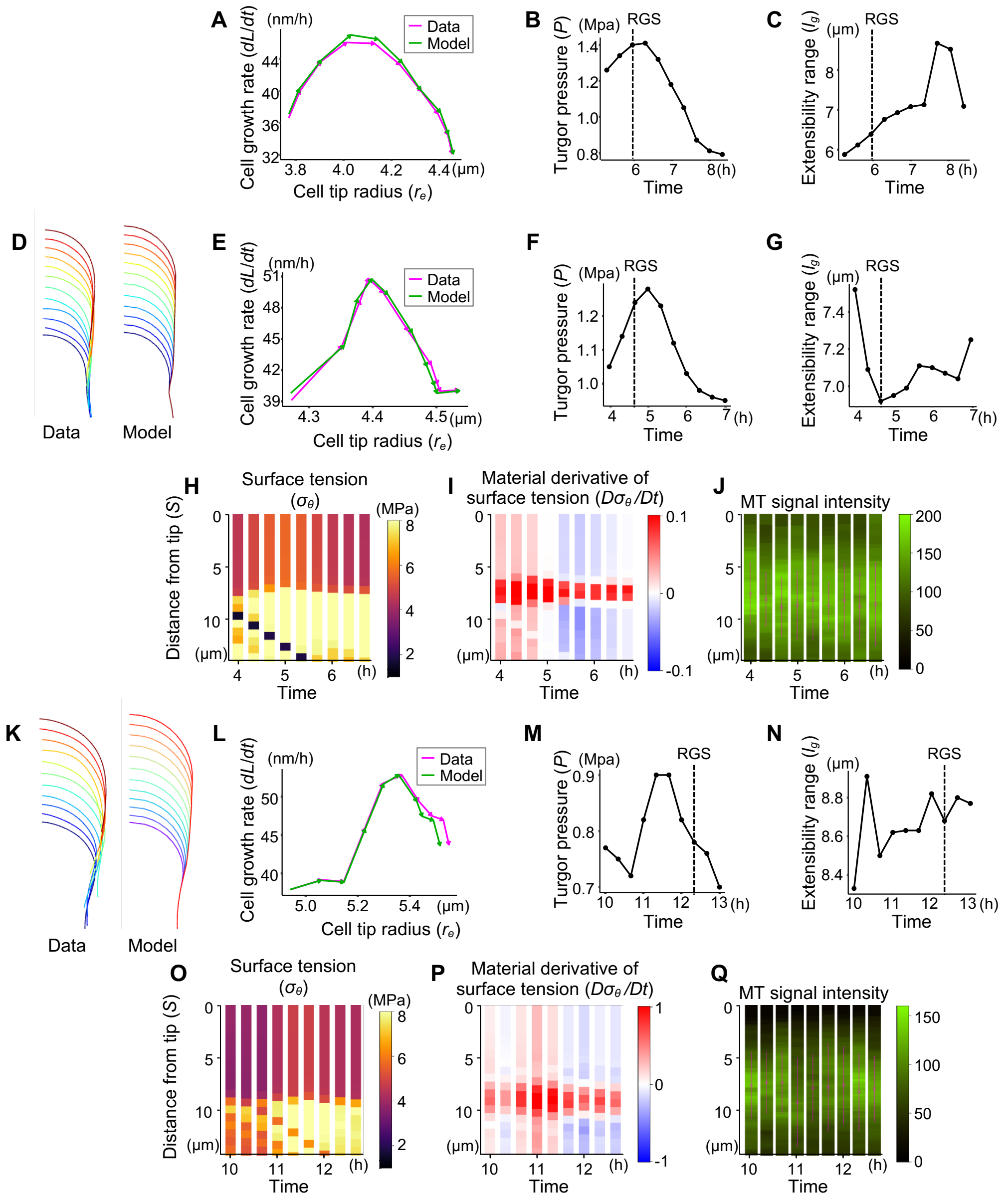


**Fig. S7. Comparison of surface tension and MT band position based on model zygote reconstruction in wild type.** (**A-C**) Results of parameter estimation for the zygote shown in Fig. 3. Timing of RGS is shown as dashed line. (**D-Q**) Results of two other representative zygotes (D-J and K-Q), analyzed as in Fig. 3. They correspond to the zygotes shown in Fig. S2A-C and S2D-F, respectively. In addition to the contour changes (D and K), comparisons of cell growth rate ($dL/dt$) and cell tip radius ($r_e$) between time-lapse observation data (Data; magenta) and reconstructed model zygote (Model; green) (A, E, L) show their nearly equal dynamics. Estimated parameters, i.e., turgor pressure ($P$) (B, F, M) and extensibility range ($l_g$) (C, G, N), show the increasement of $P$ around RGS, but no detectable correlation of $l_g$. Predicted circumferential surface tension ($\sigma_\theta$) (H and O), its material derivative ($D\sigma_\theta/Dt$) (I and P), and observed MT signal intensity (J and Q) show their correlation. Magenta lines and crosses in J and Q indicate the MT bands and their centers, respectively.

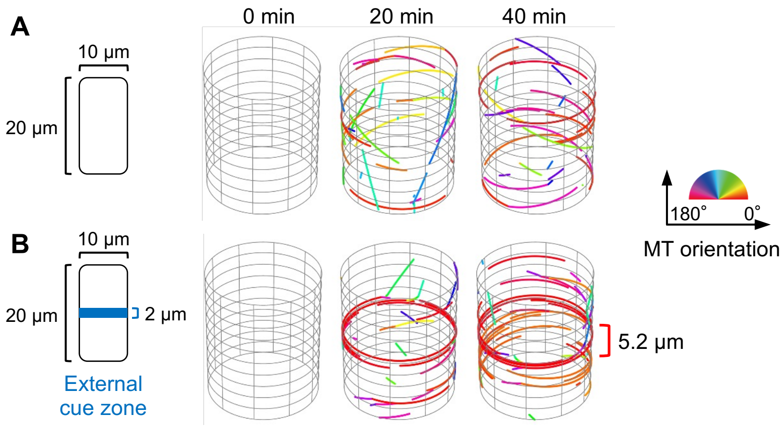


**Fig. S8. Agent-based simulation for cortical MT organization.** (**A**) Simulation results without the external directional cue using a cylindrical cell geometry mimicking the zygote (diameter 10 µm) do not exhibit MT band. For additional details, see Materials and Methods. (**B**) Simulation results with the external directional cue. Given a 2 µm wide zone that mimics a subapical region showing large material derivative, an MT band wider than the zone can be formed after 40 min. The orientation angle of the MTs is shown in color diagram.

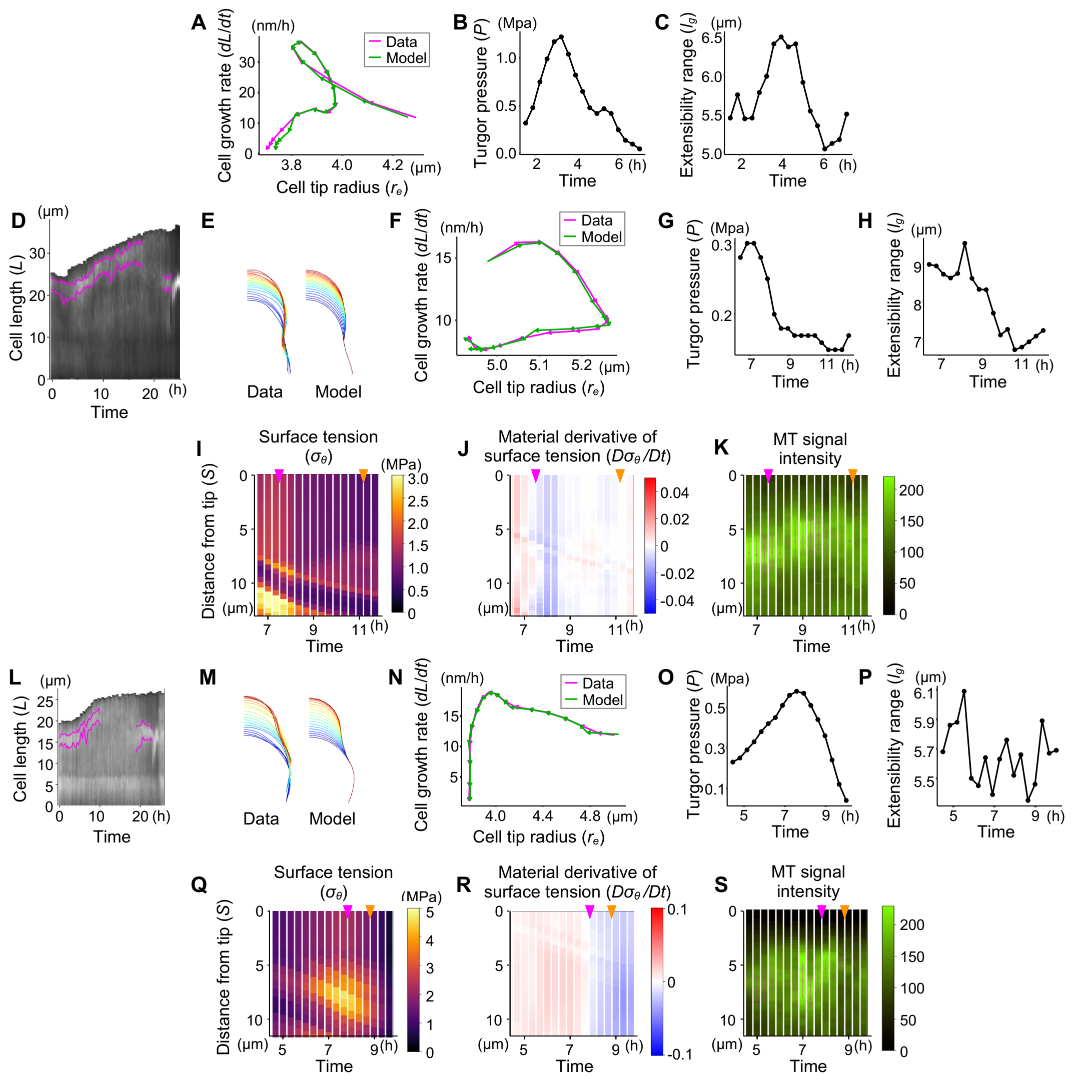

**Fig. S9. Comparison of surface tension and MT band position based on model zygote reconstruction in *yda*.** (**A-C**) Results of parameter estimation for the zygote shown in Fig. 4. (**D-S**) Results of two other representative zygotes (D-K and L-S), analyzed as in Fig. 4. In kymograph showing MT signal intensity (D and L), magenta lines indicate the upper and lower ends of the MT bands and show that PPB appearance after subapical MT band disappearance. In addition to the contour changes (E and M), comparisons of cell growth rate (*dL/dt*) and cell tip radius ($r_e$) between time-lapse observation data (Data; magenta) and reconstructed model zygote (Model; green) (A, F, N) show their nearly equal dynamics. Estimated parameters, i.e., turgor pressure (*P*) (B, G, O) and extensibility range ($l_g$) (C, H, P), show the relatively lower values of *P* compared to those of wild type shown in Fig. S7. Predicted circumferential surface tension ($\sigma_\theta$) (**I** and Q), its material derivative ($D\sigma_\theta/Dt$) (J and R) and observed MT signal intensity (K and S) show their correlation. Magenta lines and crosses in K and S indicate the MT bands and their centers, respectively. Magenta and orange arrowheads indicate when the Material derivative in surface tension is lost and when the MT bands are no longer detectable, respectively.

**Movie S1. Dynamics of cell growth and MT organization in wild type zygote.**
(Left) 2PEM image of the time-lapse observation of wild type zygote expressing MT (green)/nuclear (magenta) marker. MIP images are shown. Numbers indicate the time (hour:min) from the first frame. (Middle) MT signals are shown in white. (Right) MT signal distribution (green) that are projected onto the zygote centerline. Magenta line indicates the MT band, which is extracted by Gaussian fitting (dashed black line). Coefficient of determination ($r^2$) at each timing is shown. Scale bar: 10 μm.

**Movie S2. Dynamics cell growth and MT band disappearance in the oryzalin-treated zygote.**
(Left) 2PEM image of the time-lapse observation of oryzalin-treated zygote expressing MT (green)/nuclear (magenta) marker. MIP images are shown. Numbers indicate the time (hour:min) from the first frame. (Right) MT signals (green) that are projected onto the zygote centerline. Gaussian fitting did not work because no clear peaks were detected. Scale bar: 10 μm.

**Movie S3. Dynamics of cell growth and MT stabilization in the taxol-treated zygote.**
(Left) 2PEM image of the time-lapse observation of taxol-treated zygote expressing MT (green)/nuclear (magenta) marker. MIP images are shown. Numbers indicate the time (hour:min) from the first frame. (Middle) MT signals are shown in white. (Right) MT signal distribution (green) that are projected onto the zygote centerline. Magenta line indicates the MT band, which is extracted by Gaussian fitting (dashed black line). Coefficient of determination ($r^2$) at each timing is shown. Note that the precise MT band area could not be extracted due to distorted shape of signal peak. Scale bar: 10 μm.

**Movie S4. Comparison of surface tension and MT dynamics in wild type zygote.**
(Left) Dynamics of predicted circumferential surface tension ($\sigma_\theta$). (Middle) Material derivative of surface tension ($D\sigma_\theta/Dt$). Numbers indicate the time (hour:min) from the first frame, and values are plotted with color codes. (Right) MT signal distribution (green) that are projected onto the zygote centerline. Magenta line indicates the MT band, which is extracted by Gaussian fitting (dashed black line). Note that the right movie is a part of Movie S1.

**Movie S5. Dynamics of cell growth and MT organization in *yda* zygote.**
(Left) 2PEM image of the time-lapse observation of *yda* zygote expressing MT (green)/nuclear (magenta) marker. MIP images are shown. Numbers indicate the time (hour:min) from the first frame. (Middle) MT signals are shown in white. (Right) MT signals (green) that are projected onto the zygote centerline. Magenta line indicates the MT band, which is extracted by Gaussian fitting (dashed black line). Note that the subapical MT band disappears after 5:20 and then PPB appears at 10:40. Coefficient of determination ($r^2$) at each timing is shown. Scale bar: 10 μm.

**Movie S6. Comparison of surface tension and MT dynamics in *yda* zygote.**
(Left) Dynamics of predicted circumferential surface tension ($\sigma_\theta$). (Middle) Material derivative of surface tension ($D\sigma_\theta/Dt$). Numbers indicate the time (hour:min) from the first frame, and values are plotted with color codes. (Right) MT signal distribution (green) that are projected onto the zygote centerline. Magenta line indicates the MT band, which is extracted by Gaussian fitting (dashed black line). Note that the right movie is a part of Movie S5.

**Movie S7. Dynamics of cell growth and MT organization in *ktn1* zygote.**
(Left) 2PEM image of the time-lapse observation of *ktn1* zygote expressing MT (green)/nuclear (magenta) marker. MIP images are shown. Numbers indicate the time (hour:min) from the first frame. (Middle) MT signals are shown in white. (Right) MT signals (green) that are projected onto the zygote centerline. Magenta line indicates the MT band, which is extracted by Gaussian fitting (dashed black line). Coefficient of determination ($r^2$) at each timing is shown. Scale bar: 10 μm.

**Movie S8. Dynamics of MT organization and cell division in *trm678* zygote.**

2PEM image of the time-lapse observation of *trm678* zygote expressing MT (green)/nuclear (magenta) marker. MIP images are shown. Numbers indicate the time (hour:min) from the first frame. Scale bar: 10 μm.
